# *prolfquapp* - A User-Friendly Command-Line Tool Simplifying Differential Expression Analysis in Quantitative Proteomics

**DOI:** 10.1101/2024.10.09.617391

**Authors:** Witold E. Wolski, Jonas Grossmann, Leonardo Schwarz, Peter Leary, Can Türker, Paolo Nanni, Ralph Schlapbach, Christian Panse

## Abstract

Mass spectrometry is a cornerstone of quantitative proteomics, enabling relative protein quantification and differential expression analysis (*DEA*) of proteins. As experiments grow in complexity, involving more samples, groups, and identified proteins, traditional interactive data analysis methods become impractical. The *prolfquapp* addresses this challenge by providing a command-line interface that simplifies *DEA*, making it accessible to non-programmers and seamlessly integrating it into workflow management systems.

*Prolfquapp* streamlines data processing and result visualization by generating dynamic HTML reports that facilitate the exploration of differential expression results. These reports allow for investigating complex experiments, such as those involving repeated measurements and multiple explanatory variables. Additionally, *prolfquapp* supports various output formats, including XLSX files, SummarizedExperiment objects and rank files, for further interactive analysis using spreadsheet software, the *exploreDE* Shiny application, or gene set enrichment analysis software.

By leveraging advanced statistical models from the prolfqua R package, *prolfquapp* offers a user-friendly, integrated solution for large-scale quantitative proteomics studies, combining efficient data processing with insightful, publication-ready outputs.

**TOC Graphic:** 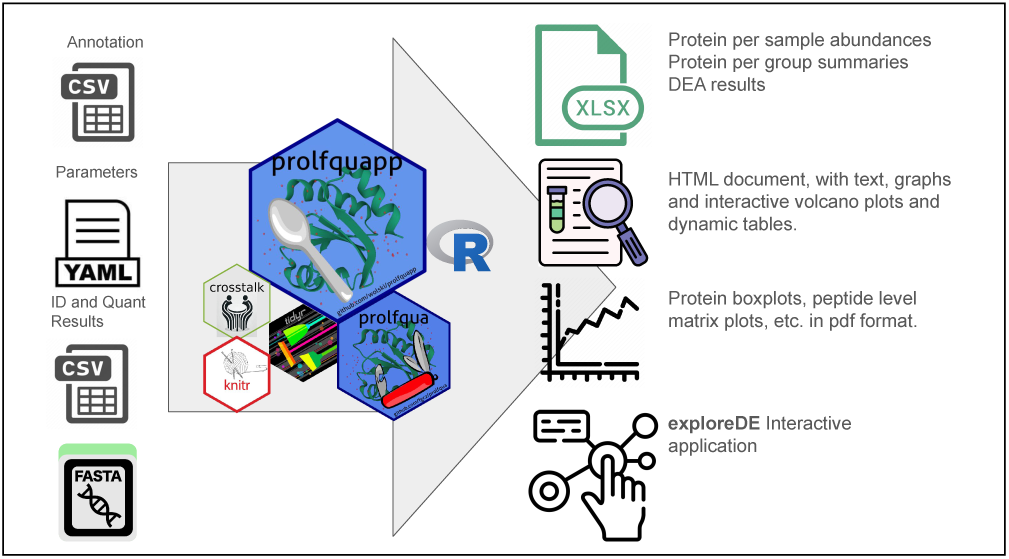

This visual table of contents illustrates the workflow and key features of the *prolfquapp* tool for differential expression analysis in proteomics. On the left are the inputs, like the CSV for annotation and quantification results, YAML for parameters, and FASTA files for protein information. In the center are the *prolfquapp* and *prolfqua* R packages and supporting tools like crosstalk and knitr, representing the core processing components. On the right side, the figure highlights the various outputs generated by *prolfquapp*

1. XLSX files containing protein abundances, group summaries, and differential expression results.
2. HTML reports with text, graphs, interactive volcano plots, and dynamic tables for data exploration.
3. PDF documents with detailed protein boxplots and peptide-level matrix plots.
4. Integration with *exploreDE* for interactive data visualization.

This diagram concisely summarizes the flow from data input to the creation of analysis-ready outputs, offering a clear overview of the prolfquapp toolset.

## Introduction

The *prolfqua* R package^1^ for differential expression analysis is becoming increasingly popular and has been used in several recently published proteomics studies^2–5^. The prolfquapp package, which builds upon the *prolfqua*package, implements a ready-to-use command line application that starts from outputs of popular quantification software ^6–8^, and allows to analyze experimental designs most commonly used for protein expression analysis ^9,10^. While the *prolfqua* package is a tool for bioinformaticians, prolfquapp’s primary users are researchers conducting quantitative proteomics studies.

*Prolfquapp* enables users to execute a differential expression analysis without knowing a programming language and simplifies the integration into workflow managers. The software performs protein or peptide differential expression analysis and generates dynamic HTML reports that contain quality control plots and interactive visualizations. Furthermore, we export results in multiple formats, including XLSX files, .rnk or .txt files for gene set enrichment and overrepresentation analysis, and the SummarizedExperiment^11^ for interactive visualization in the *exploreDE* ^12^ application.

*Prolfquapp* simplifies and automates quality control and differential expression analysis, of quantitative proteomics data. Other R packages, such as *Einprot* ^13^, *LFQAnalyst* ^14^, *MS-Dap* ^15^, *Amica* ^16^, or *MSstatsShiny* ^17^ have a similar aim, to make *DEA* analysis end user friendly. To model the differential expression analysis, or to perform the data preprocessing, these packages use methods implemented in the packages *limma* ^18^, *MSstats*, or *msnbase* ^19^, while we use models and methods implemented in the package *prolfqua* ^1^. These R packages enable non-programmers to perform *DEA* by implementing a graphical user interface using *Shiny* ^14,17,20,21^, or implement a facade^22^ that provides a simplified interface to otherwise complex code and allows running a complex analysis with only a few lines of code.^13,15,17^ However, for large-scale experiments, when reading the data, and model fitting takes a long time, these packages might not be ideal.

The *prolfquapp* package advocates separating the model fitting from the interactive analysis of the results. We furthermore recommend delaying the data filtering for missing values or the number of peptides identified after modeling the differential expression. The rationale is that keeping measurements with missing values does not relevantly affect the normalization, the p-value moderation, or the FDR estimation. In the worst case, we make the FDR estimates more conservative by performing the p-value adjustment on a larger number of tests. Therefore, we estimate the fold changes of as many proteins and peptides as possible and recommend filtering the results. Another benefit is that we do not need to rerun the relatively time-consuming model fitting. To facilitate this approach, in addition to the differential expression results, we provide summary statistics for each protein and peptide, such as the number of peptides per protein in the experiment or sample or the number of missing values per group. Based on this information, we can refine the lists of significant proteins or the list of fold changes to include in the over-representation analysis (ORA) or incorporate them into gene-set enrichment analysis (GSEA), respectively. This filtering can be performed interactively using spreadsheet software such as Excel or OpenOffice since we provide the results in XLSX files. Also, when using the *exploreDE* application, which reads SummarizedExperiment (SE) output generated by the *prolfquapp*, we can filter the data interactively by the number of peptides per protein observed in the experiment.

Data management systems such as B-Fabric^23^ and OpenBIS,^24^ which schedule bioinformatics compute jobs via platforms like Galaxy^25^ and SLURM,^26^ utilizing tools such as FragPipe and DIA-NN, and enable reproducible science, are critical in modern bioinformatics. These systems automate complex analyses and manage large datasets, enhancing efficiency and scalability. We developed *prolfquapp* to integrate seamlessly with these platforms, enabling users to efficiently queue differential expression analysis (*DEA*) jobs while ensuring reproducibility across diverse computational environments. With its command-line interface, *prolfquapp* can be easily incorporated into these workflow management systems, increasing its utility for large-scale experiments where interactive analysis may be impractical. Furthermore, we wanted to enable our users, frequently using Windows computers, to replicate or modify the analysis by updating the annotation and parameters.

## Methods

### Sample Annotation

*Prolfquapp* formalizes how to annotate samples with explanatory variables for some of the most common experimental designs: parallel group design and factorial designs with and without repeated measurements. We defined an annotation file format that provides the explanatory variables and specifies the group comparisons. The first column is reserved for the sample identifier, which is either the raw file name in the case of label-free experiments or the channel for labeled experiments. The values provided in that column must match the sample names in the quantification software outputs. We support the creation of annotation files using the prolfqua_dataset script which extracts the sample names used by the quantification software. The column names can be upper or lowercase. Further columns are used to assign the explanatory variables and define the contrasts to compute (see table 1). The column *subject*, which can be used to provide, for instance, patient identifiers in paired experiments, is optional. Instead of the column *control*, the columns *ContrastName* and *Contrast* can be provided to specify the differences to compute (see Table 4).

**Table 1:** Specification of the table we use to annotate the samples and specify the experimental design, as well as contrasts. We compare all groups to the control reference group specified in the control column.

| Column Name | What |
| --- | --- |
| relative.path/path/raw.file/channel | The sample identifier for each row must be unique |
| name | Sample name used in tables and figures |
| group/experiment | Main factor |
| subject/bioreplicate | Blocking factor (optional) |
| control | Specify the control (C) or treatment (T) condition |

### Integrating Protein Information from FASTA files

Because there are differences in what information about proteins is provided in the outputs of the quantification software, *prolfquapp* enriches the quantification data with protein information extracted directly from the FASTA file. We parse the FASTA file using the *seqinr* R package^27^ to compute protein lengths and the number of theoretical tryptic peptides, use them to compute intensity-based Absolute Quantitation (iBAQ) values, and parse the protein descriptions to extract gene names. We are adding this information to generated reports.

### Parsing of Quantification Results

Developing an application that integrates multiple upstream processing software presents challenges in handling diverse tabular file formats. These challenges go beyond column naming or format (tidy vs. wide) and include differences in how quantification software extracts protein identifiers from FASTA files. Different tools may assume UniProt format or interpret identifiers in varying ways, sometimes leading to unpredictable behavior. Additionally, quantification files may vary regarding the sequences included (forward only or with decoys) and the filtering based on the q-value thresholds.

There are also inconsistencies in how quantification software represents file names, with some tools removing extensions, others retaining them, and some even keeping the full file path, which may differ based on the operating system. Despite efforts to standardize file structures across software versions, import functions in tools like *prolfqua* often require updates to accommodate variations in quantification software, operating systems, and FASTA file formats such as UniProt or Ensembl.

*Prolfquapp* builds on the *prolfqua* package, which stores all the data in a tidy data table containing the explanatory and response variables for all the proteins. A configuration object provides annotations of the columns in the tidy data table. Therefore, when using *prolfqua* and *prolfquapp*, the crucial task is correctly parametrizing the configuration object. We automate the process of parsing quantification results and creating a configuration for various output formats. Table 2 shows which output formats are currently supported and if protein or peptide-level *DEA* is possible.

**Table 2:** Quantification software supported at the time of publishing. The second column shows which file is used as input. The third column shows which parameter must be passed to the prolfqua_dea and prolfqua_qc applications to generate protein or peptide centric repots.

| Software | Input file | –software |
| --- | --- | --- |
| DIANN/FragPipe DIA | report.tsv | **DIANN** or **DIANN_PEPTIDE** |
| FragPipe DDA workflow | msstats.tsv | **MSSTATS** or **MSSTATS_PEPTIDE** |
| FragPipe TMT workflow | PSM.tsv | **FP_TMT** or **FP_TMT_PEPTIDE** |
| MaxQuant | peptide.txt | **MAXQUANT** or **MAXQUANT_PEPTIDE** |
| Spectronaut | BGS report | **BGS** or **BGS_PEPTIDE** |

Although we provide implementations for some quantification software, we also make it possible to inject custom file-parsing functions from other R packages if they are compatible with the *prolfquapp* parsing interface. The *prolfquappPTMreaders* package^28^ is an example of an external package that implements *prolfquapp*-compatible methods for reading outputs of FragPipe PTM workflows.

### Design and Parameter Management

We pass parameters to the programs using command line arguments or YAML configuration files. YAML is structured, easy to read, and allows users to define key parameters, such as the data normalization method. It also includes details about the input data, project, and work units, which we integrate into the HTML reports. While this may not be crucial for single datasets, it becomes vital when managing multiple datasets or interoperating with a LIMS system.

To modularize and structure the code, we utilized R6 classes. For instance, the Annotation-Processor class handles methods for parsing the sample annotation file, AppConfiguration manages the YAML parsing, QCGenerator contains functionality for the QC application, and DEAAnalyse organizes methods for the *DEA* application.

### Generating interactive HTML and XLSX documents

An important functionality of the *prolfquapp* package is to report *DEA* results. We use the R bookdown package^29^ to generate the main HTML report, which allows us to create figure and table captions and reference them in the text. We used *ggplot2* ^30^ and *plotly* ^31^ to create figures and add interactive features such as hovering and highlighting. We implement tables using the *DT* package^32^, enabling sorting, searching, and filtering of data. The tables interact with the plots through the crosstalk^33^ package, allowing us to highlight selected items in the plots based on table selections for linked exploration. Additionally, we use the *writexl* package^34^ to generate XLSX files that include protein intensity estimates and statistical results for further analysis.

## Results and Discussion

### Workflow

To quantify peptidoform or protein abundances, we use protein quantification software that processes mass spectrometry raw data and FASTA files. The quantification software stores the results in tabular text files within a directory. Here we show how the *prolfquapp* command line tools can be executed on a Linux or macOS shell to generate QC reports and perform a DE analysis (for Windows, we provide bat files).

The first step when modeling the peptide and protein abundances is to generate a file that annotates the samples with explanatory variables. Ideally, this information is stored in a LIMS system, and using an API, the data can be extracted and written into an annotation file. In case there is no LIMS system, we can run the command:

`./prolfqua_dataset.sh -i data_dir/ -s DIANN -d annotation.xlsx`

, which will parse the quantification result from the data_dir and extract the names of all raw files (label-free) or channels (labeled quantification) and prepare a table to add sample names, the grouping variable, and contrast information (Table 1, Table 2).

The next step is to run the quality control, which generates HTML outputs that visualize the data and help to identify outlier samples or determine if samples were contaminated. Table 5 summarizes outputs generated by the QC application.

`./prolfqua_qc.sh -i data_dir/ -d annotation.xlsx -s DIANN -o results_dir`

If the QC shows outlier samples, e.g., samples with fewer peptides and proteins or samples that unexpectedly cluster with samples from a different group, or if there is a noticeable batch effect, we would edit the annotation file. For instance, we could remove rows with outlier samples or add an explanatory variable using the column *subject* to model the batch effect (see Table 3).

**Table 3:** Annotation file listing a subset of raw files, sample names, sample groups (Group), and patient identifiers (Subject). The values from the “Name” column will be used in the report figures and tables; therefore, they should be short yet descriptive.

| raw.file | Name | Group | Subject |
| --- | --- | --- | --- |
| CPTAC_CCRCC_W_JHU_20190112_LUMOS_C3L-00004_NAT | NAT_Old_4 | NAT_Old | 4 |
| CPTAC_CCRCC_W_JHU_20190112_LUMOS_C3L-00004_T | T_Old_4 | T_Old | 4 |
| CPTAC_CCRCC_W_JHU_20190112_LUMOS_C3L-00010_NAT | NAT_Young_10 | NAT_Young | 10 |
| CPTAC_CCRCC_W_JHU_20190112_LUMOS_C3L-00010_T | T_Young_10 | T_Young | 10 |
| CPTAC_CCRCC_W_JHU_20190112_LUMOS_C3L-00011_NAT | NAT_Old_11 | NAT_Old | 11 |
| CPTAC_CCRCC_W_JHU_20190112_LUMOS_C3L-00011_T | T_Old_11 | T_Old | 11 |
| CPTAC_CCRCC_W_JHU_20190112_LUMOS_C3L-00026_NAT | NAT_Old_26 | NAT_Old | 26 |
| CPTAC_CCRCC_W_JHU_20190112_LUMOS_C3L-00026_T | T_Old_26 | T_Old | 26 |
| CPTAC_CCRCC_W_JHU_20190112_LUMOS_C3L-00079_NAT | NAT_Young_79 | NAT_Young | 79 |

A configuration file with information about the experiment and analysis parameters is needed to run the differential expression analysis. Here, we specify, for instance, the normalization method, the protein intensity estimation method, or thresholds used to visualize significant fold changes. We create the configuration with default settings by calling the command: ./prolfqua_yaml.sh config.yaml

Finally, the *DEA* analysis can be run by calling the command:

`./prolfqua_dea.sh -i data_dir/ -d annotation.xlsx -y config.yaml -s DIANN`

In a workflow management setting, the annotation and YAML files are generated based on information stored in the LIMS system. For B-Fabric, we create the annotation and YAML files using the *bfabricPy* library^35^.

### Analyzing an experiment with two factors

To demonstrate how we use *prolfquapp* to analyze a dataset with two factors and repeated measurements, we used the renal cell carcinoma dataset^36^ as an example. In this study treatment-naive tumors (T) and paired normal adjacent tissues (NAT) were collected in young and old patients. Table 1 shows the annotation file. It is worth noting that although this experiment has two factors, cell type, and age, we merge them, and the model uses a single explanatory variable, Group. In addition, we specify the patient identifier in the column Subject. We specified the model in this way because we were interested in looking into the interaction between age and cell type, and furthermore, we wanted to block for between patient differences.

Finally, we need to specify the contrasts, which we do by adding two more columns to the annotation file, one containing the name of the comparison (ContrastName) and the other a formula expressing the differences (Contrast). The first contrast T_vs_NAT examines the differences in tumor and adjacent tissue; the contrast Old_vs_Young, the difference between old and young; the Contrasts T_vs_Nat_gv_Young and T_vs_NAT_gv_Old the differences within the groups of young and old patients, while the last examines if this difference depends on the age of the patients (DoesTvsNatDependsOnAge). The full analysis is available to review the generated outputs from the ’*DEA* Large Dataset Example’ web resource^37^.

**Table 4:** Table showing two additional columns used to specify the contrasts. Please note that the group names have the prefix “G_” (shorthand for Group).

| ContrastName | Contrast |
| --- | --- |
| T_vs_NAT | $(G\_T\_Old + G\_T\_Young)/2 - (G\_NAT\_Old + G\_NAT\_Young)/2$ |
| Old_vs_Young | $(G\_NAT\_Old + G\_T\_Old)/2 - (G\_NAT\_Young + G\_T\_Young)/2$ |
| T_vs_NAT_gv_Young | $G\_T\_Young - G\_NAT\_Young$ |
| T_vs_NAT_gv_Old | $G\_T\_Old - G\_NAT\_Old$ |
| DoesTvsNatDependsOnAge | $T\_vs\_NAT\_gv\_Old - T\_vs\_NAT\_gv\_Young$ |

### Outputs - interactive HTML report and support of downstream analysis

The *prolfquapp* provides a comprehensive set of outputs, including interactive HTML reports, XLSX files, or serialized R objects, to facilitate data interpretation. Table 5 lists the output files generated by the QC and DEA applications.

**Table 5:** List of output files generated by *prolfquapp* applications for QC and DEA (see Supporting Material 1).

| Application | File name | File content |
| --- | --- | --- |
| prolfqua_qc | QC_sampleSizeEstimation.html | Experiment QC and Sample Size Estimation Report |
|  | QC_porteinAbundance.html | Interactive HTML report displaying protein iBAQ values |
|  | proteinAbundances.xlsx | Spreadsheet with peptide and protein intensity estimates, protein iBAQ values, and protein statistics per group (CV, nr of missing) |
| prolfqua_dea | DE_.html | DE Analysis Report, with e.g., interactive volcano plots |
|  | QC_*.html | DE Quality Control Report |
|  | ORA_*.txt | Files for all contrasts to perform ORA analysis |
|  | GSEA_*.rnk | Files for all contrasts to perform GSEA analysis |
|  | DE_*.xlsx | Spreadsheet file with protein abundance estimates, differential expression results, and additional summaries and statistics |
|  | IBAQ_*.xlsx | Spreadsheet file with iBAQ protein abundances |
|  | SummarisedExperiment.rds | Serialized SE object used by the *exploreDE* application |

The QC application generates a QC and sample size estimation report and an interactive protein or peptide abundance report (Figure 2). Furthermore, we also write an XLSX file containing either peptide abundances, or protein abundance estimates determined using the Tukeys median polish, protein *iBAQ* values, and summaries such as the number of peptides per protein and experiment or sample. The protein quantity estimates are an alternative to the protein-level summaries produced by the quantification software, that typically reports *maxLFQ* abundances. Furthermore, using the values in the XLSX file, we can reproduce the figures in the QC report.

**Figure 1:**
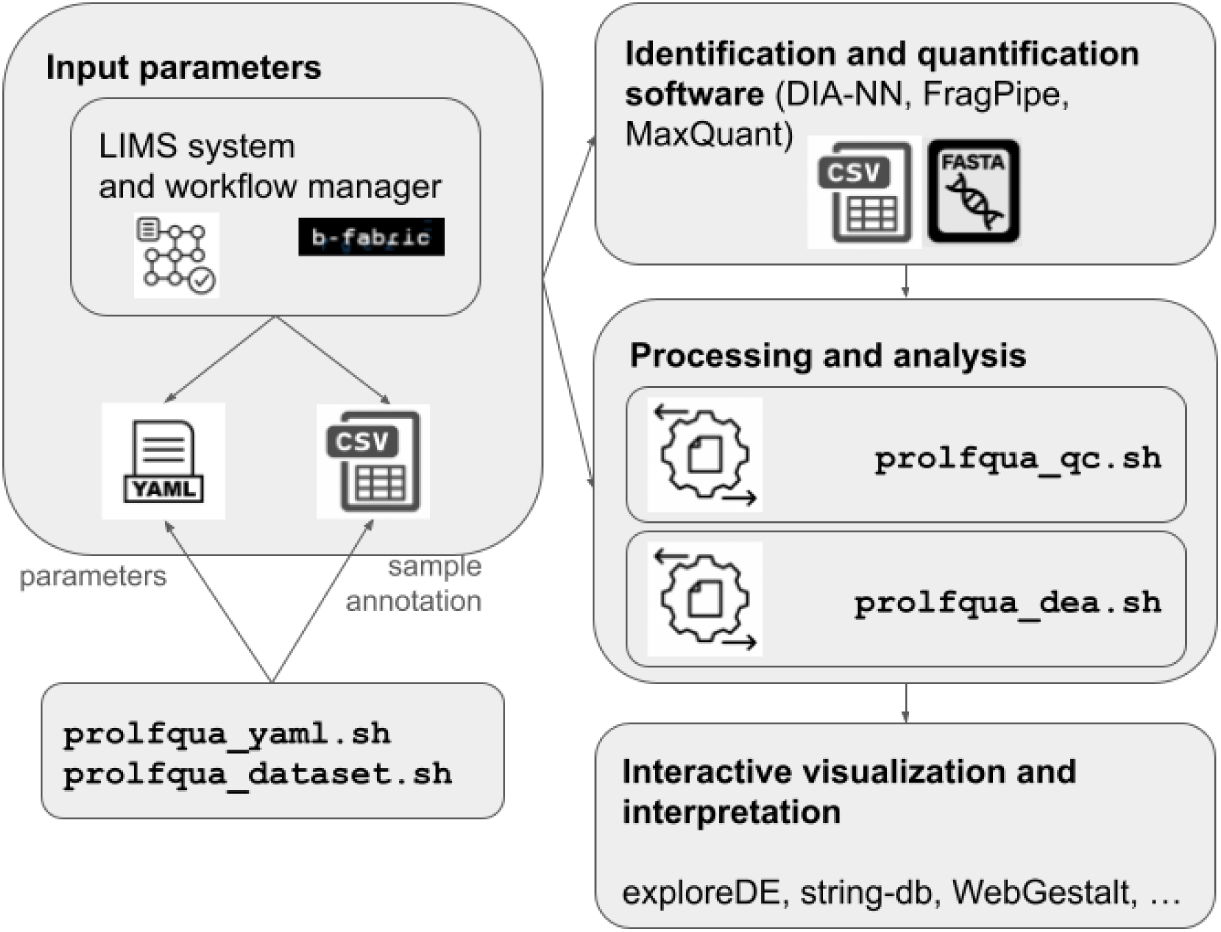
The *DEA* analysis workflow using *prolfquapp*. Input parameters and sample annotation are either generated by the LIMS system or a workflow manager, or using the prolfqua_yaml and prolfqua_dataset scripts. Then, the prolfqua_qc and prolfqua_dea scripts can be run to analyze the identification and quantification software outputs. We support interactive data visualization and data interpretation by generating outputs compatible with *exploreDE* and *string-db*.

**Figure 2:**
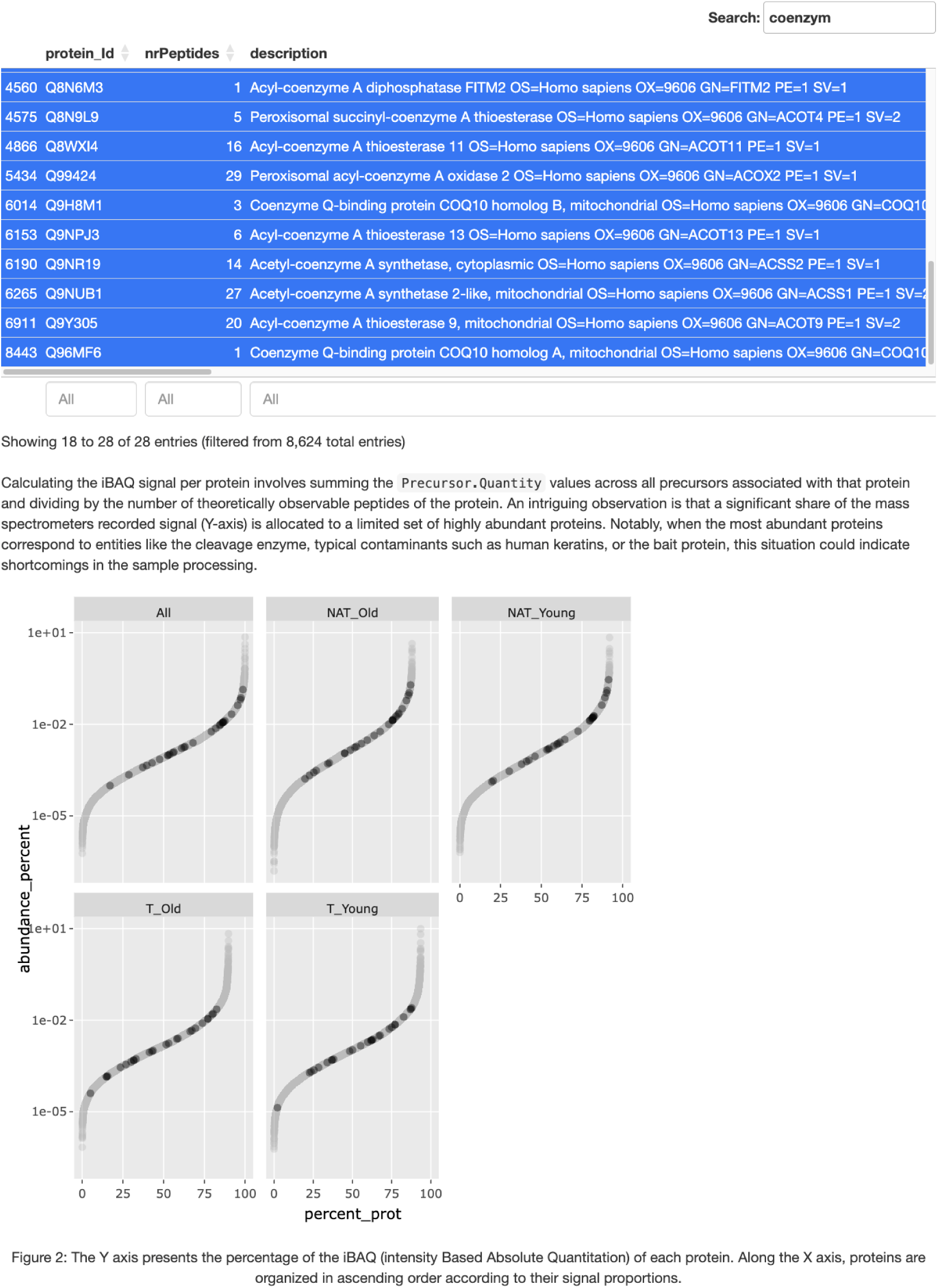
Screenshot of the interactive abundance visualization in the QC HTML report. The plot shows the protein iBAQ values for each group. The plot is interactive, allowing users to select proteins, for instance, all having the word coenzyme in the description line and highlighting them in the figure.

The *prolfquapp DEA* application generates a folder with two subfolders, one containing all the input files needed to replicate the analysis and a folder containing all the result files. The resulting folder can be archived, and the data in the input subfolder can be used to replicate or, by changing parameters or editing the annotation file, modify the analysis.

The primary output is the report in HTML format. The document starts by introducing differential expression analysis. Then, we show visual summaries of protein and peptide identification, such as the number of identified proteins or the distribution of missing observations. The report visualizes the results of the *DEA* using volcano plots for all studied contrasts. These volcano plots are interactive and linked when there are multiple comparisons. This functionality allows users to highlight a protein across all volcano plots by clicking on a data point, making it easier to identify quantitative and qualitative interactions, or search for proteins of interest in the table with the *DEA* results (see Figure 3). The report also includes dynamic tables that allow users to search and filter proteins of interest, which can be highlighted in the volcano plot. Next, we summarize the numbers of differentially expressed proteins and visualize them using an upset plot, which displays the intersections of significant proteins or peptides between multiple sets of contrasts (see Supporting Material 1). The section dedicated to additional analysis provides pathways for downstream omics analysis, such as gene set enrichment analysis, with direct links to web services like WebGestalt and the String functional protein association network.

**Figure 3:**
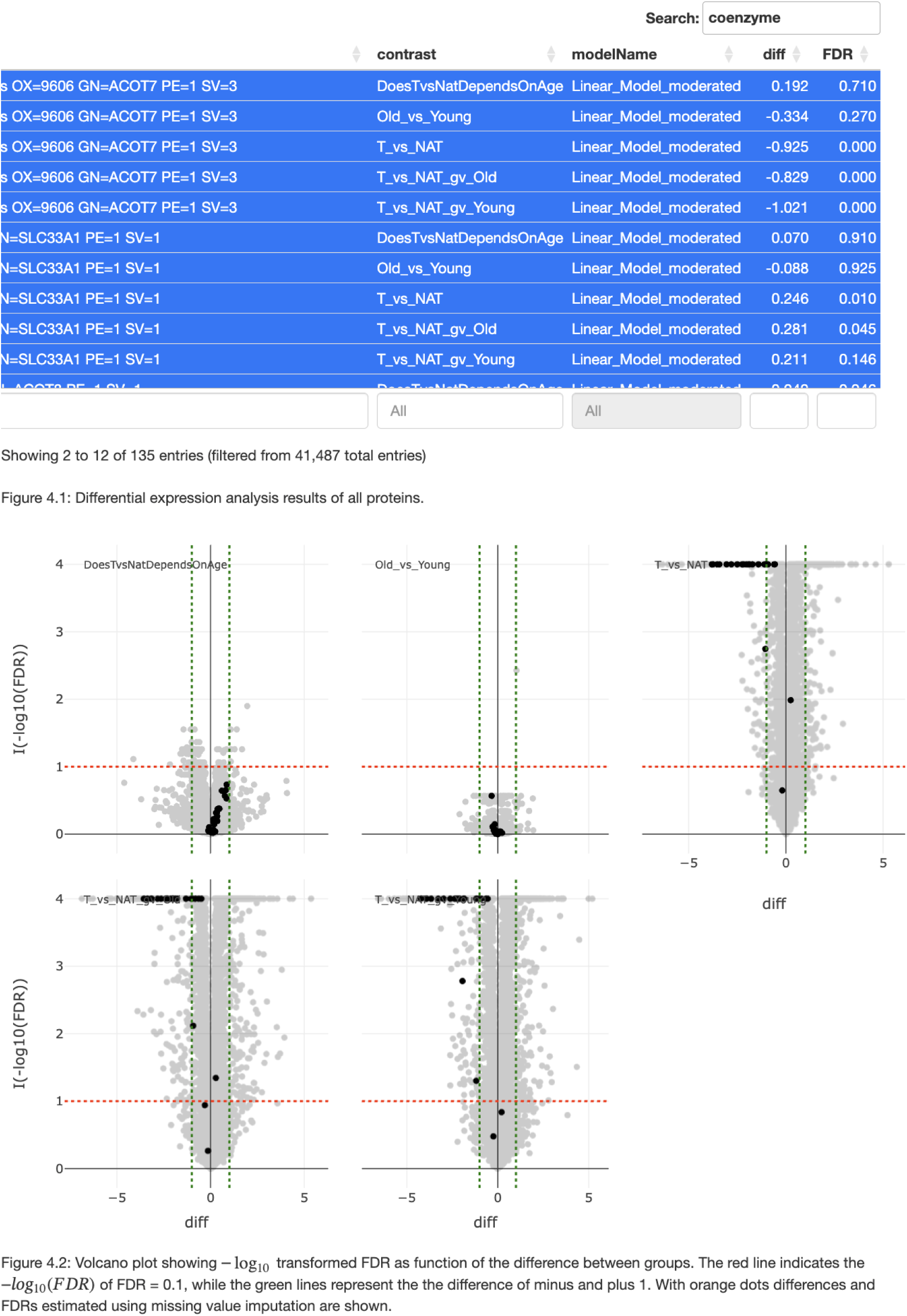
Interactive volcano plots in the DEA HTML report. The plot is interactive, allowing users to select proteins, for instance, all having the word coenzyme in the description line and highlighting them in the figure.

In addition, we provided a *DEA*-QC HTML document with plots and tables, which we found informative but did not make it into the main report. The figures there, for instance, help to asses if the data normalization method was able to reduce the standard deviation of proteins within groups compared to only log2 transformed intensities (see Supporting Material 1). This document also contains histograms showing the distribution of the p-values for each contrast, an important plot helping to detect potential issues in a dataset, such as biases or errors in the statistical model.

While the HTML report offers an interactive and visual exploration of the results, the main output of *prolfquapp* is an XLSX file. The XLSX file includes comprehensive tables with peptide and protein intensity estimates, IBAQ values, differential expression analysis results, and various protein or peptide summaries. These summaries provide essential metrics, such as the number of peptides per protein, peptide counts per sample, and protein coefficient of variation (CV), allowing for the filtering of protein coverage, proportion of missing values, or highly variable proteins. Spreadsheet programs remain a widely used tool among life science researchers and are ideal for filtering the results. Furthermore, figures and plots from the HTML report can be fully reproduced using the data contained in the XLSX output.

A conscious design choice was not to include, downstream analysis such as ORA or GSEA, or functionality to change colors in figures or figure layouts. The reason here is that there are excellent tools for ORA and GSEA such as *WebGestalt* ^38^, *MSigDB* ^39^, or *string-db* ^40^ that can perform downstream analysis of omics data. Furthermore, we do want to run the application without user interactions and therefore we generate SE files compatible with *exploreDE*.

With its dual approach of offering both interactive HTML reports and data-rich XLSX files, *prolfquapp* is a versatile tool that can adapt to the needs of both exploratory and detailed analytical workflows.

### Interactive visualization using exploreDE

The importance of interactively visualizing the data, especially the ability to easily create publication-ready visualizations by life science experts, must be considered. Therefore, the seamless integration of *prolfquapp* with the *exploreDE* application is an essential cornerstone of the differential expression analysis pipeline. The *exploreDE* application allows the creation of highly customizable heatmaps, volcano, or PCA plots, which can be exported in various formats and included in presentations, reports, and publications. The Supporting Material 2^41^ shows the interactive visualization of the renal cell carcinoma dataset using the *exploreDE* application.

### Reproducibility and Replicability

*Prolfquapp* enhances reproducibility and replicability by generating self-contained outputs encapsulating all necessary parameters and settings. Lastly, it is essential to report all the steps performed, the execution progress, and errors for non-interactive applications. For this task, we use the R *logger* library^42^.

To simplify running *prolfquapp* and enhance reproducibility, we provide container images alongside the R package. If Docker or Podman is installed on your system, a script^43^ available in the repository will handle the setup and execution of the container, which allows users without experience in container technology to execute any of the command-line scripts within a versioned environment.

### Performance

The renal cell carcinoma dataset comprises 187 samples, with 8613 proteins identified using the FragPipe DIA-NN workflow. Running the *DEA* with the prolfqua_dea took approximately 20 minutes and took 20GB of RAM on an Apple M1 MacBook Pro with 32GB of RAM.

### Software Availability

The *prolfquapp* package is available as open-source software and can be accessed via github repository^44^. The package is compatible with Windows, macOS, and Linux operating systems. Instructions for installation, usage, and documentation are provided in the repository. The software is released under the MIT License, allowing free use, modification, and distribution. We encourage the community to report any issues or bugs via the GitHub issue tracker and welcome contributions in the form of code submissions or feature requests.

### Conclusion

Looking ahead, we are committed to expanding *prolfquapp*’s functionality to meet the evolving needs in proteomics data analysis. One planned feature is integrating the *prozor* R package^45^ to provide an additional option for determining peptide-to-protein relationships, offering users more flexibility in their analysis. We also plan to report peptide-level fold changes in addition to the protein-level fold changes by default, providing deeper insights and a more detailed understanding of differential expression results. Furthermore, we aim to offer more options to improve the modeling of missing observations and include count-based models.

The *prolfquapp DEA* application creates organized folders containing all relevant inputs (annotation, parameters in YAML file) and outputs (e.g., results, reports), preserving all analysis components. Additionally, we provide Docker images to guarantee a consistent computational environment, allowing users to replicate the exact setup. In this way, *prolfquapp* helps scientists meet the requirements of funding agencies, journals, and academic institutions while publishing their data according to the FAIR data principles.

We designed *prolfquapp* so we can integrate it into scalable, high-throughput, and reproducible workflows by streamlining the configuration and execution. Its seamless integration with workflow management systems makes it particularly suitable for large-scale experiments where interactive analysis is impractical. By providing detailed, informative reports and interactive visualization of differential expression analysis results, *prolfquapp* lowers the barrier for end users to engage in advanced proteomics analysis. Furthermore, the integration with the *exploreDE*application allows the generation of publication-ready visualizations, giving the end users the independence to present their results. Through this, we made *prolfquapp* a practical solution for a wide audience within the proteomics community. Looking ahead, we anticipate that community contributions, such as bug reports and feature suggestions, will play an important role in shaping future versions of *prolfquapp*, helping ensure that it continues to meet the needs of the field as it evolves.

## Abbreviations

API: application programming interface
CV: Coefficient of Variation
DEA: Differential Expression Analysis
DIA: data independent acquisition
FAIR: Findable, Accessible, Interoperable, and Reusable
GSEA: Gene Set Enrichment Analysis
iBAQ: intensity-based Absolute Quantification
LC: Liquid Chromatography
LC-MS: Liquid Chromatography followed by Mass Spectrometry
MS: mass spectrometry
ORA: Over-Representation Analysis
PCA: Principal Component Analysis
QC: Quality Control
SE: SummarizedExperiment
TMT: Tandem Mass Tag

## Supporting Information

- Supporting Material S1: Example analysis of a dataset with two factors^37^

– docker.sh file is used to set up the docker container and run the analysis by executing /dockers.sh quarto render Readme.qmd
– README.html shows the code to execute the analysis
– folder starting with FragPipe_ contain the inputs
– folder starting with QC_ contain the output of the prolfqua_qc application
– folder starting with DEA_ contain the output of the prolfqua_dea application

Supporting Material S2: *Prolfquapp DEA SummarizedExperiment* visualized with *exploreDE* ^41^

## Acknowledgement

The authors would like to thank the staff and community of the Functional Genomics Center Zurich (FGCZ) for their continued support, collaboration, and valuable discussions. Special appreciation goes to Laura Kunz, Bernd Roschitzki, Sibylle Pfammatter, Tobias Kockmann, and Alaa Othman for their critical input and contributions.

